# Temporal dynamics of Na/K pump mediated memory traces: insights from conductance-based models of *Drosophila* neurons

**DOI:** 10.1101/2023.03.07.531036

**Authors:** Obinna F. Megwa, Leila May Pascual, Cengiz Günay, Stefan R. Pulver, Astrid A. Prinz

## Abstract

Sodium potassium ATPases (Na/K pumps) mediate long-lasting, dynamic cellular memories that can last tens of seconds. The mechanisms controlling the dynamics of this type of cellular memory are not well understood and can be counterintuitive. Here, we use computational modeling to examine how Na/K pumps and the ion concentration dynamics they influence shape cellular excitability. In a *Drosophila* larval motor neuron model, we incorporate a Na/K pump, a dynamic intracellular Na^+^ concentration, and a dynamic Na^+^ reversal potential. We probe neuronal excitability with a variety of stimuli, including step currents, ramp currents, and zap currents, then monitor the sub- and suprathreshold voltage responses on a range of time scales. We find that the interactions of a Na^+^-dependent pump current with a dynamic Na^+^ concentration and reversal potential endow the neuron with rich response properties that are absent when the role of the pump is reduced to the maintenance of constant ion concentration gradients. In particular, these dynamic pump-Na^+^ interactions contribute to spike rate adaptation and result in long-lasting excitability changes after spiking and even after sub-threshold voltage fluctuations on multiple time scales. We further show that modulation of pump properties can profoundly alter a neuron’s spontaneous activity and response to stimuli by providing a mechanism for bursting oscillations. Our work has implications for experimental studies and computational modeling of the role of Na/K pumps in neuronal activity, information processing in neural circuits and the neural control of animal behavior.

## Introduction

Sodium potassium ATPases (Na/K pumps) are ubiquitous components of neuronal membranes. They use ATP to pump Na^+^ out of neurons and K^+^ into neurons against their respective concentration gradients, and play a critical role in maintaining the resting membrane potential of neurons (reviewed in Kaplan, 2002). There is a growing realization that in addition to playing this housekeeping role, Na/K pumps also play surprisingly dynamic roles in regulating cellular excitability. In particular, Na/K pumps mediate long-lasting membrane afterhyperpolarizations (AHPs) in response to stimulus-induced high spiking activity. These AHPs can provide a neuronal memory of previous activity than can last tens of seconds (reviewed in Picton et al., 2017b). Pump mediated AHPs were first characterized in invertebrate neurons (Scuri et al., 2007, Pulver and Griffith, 2010) but have now been found in *Xenopus* (Zhang and Sillar, 2012) and mouse spinal neurons (Picton et al., 2017a).

In all species studied to date, the fundamental mechanisms of Na/K pump mediated AHPs appear remarkably conserved. High frequency bouts of spiking in neurons trigger long lasting outward currents with little or no conductance changes for tens of seconds. These long lasting hyperpolarizations are subject to neuromodulation (Picton et al., 2017a, Hachoumi et al., 2022) and interact with voltage gated ionic conductances to shape neuronal intrinsic properties (Picton et al., 2018, Pulver and Griffith, 2010). Despite this growing awareness of Na/K pumps as dynamic players in tuning cellular excitability within neural circuit components, our understanding of how specific Na/K pump properties shape intrinsic properties generally and the relationships between spiking and AHP properties remains relatively fragmented. This is in part due to a lack of pharmacological or genetic tools for precisely manipulating specific pump properties and difficulty of experimentally controlling and monitoring intra- and extracellular Na^+^ and K^+^ concentrations while also maintaining physiologically realistic conditions for the neurons under study.

Neuroscience researchers have increasingly turned to computational modelling of Na/K pumps to circumvent challenges associated with studying pumps in experimental preparations. In multiple model systems, Hodgkin-Huxley-type model neurons that incorporate both voltage-gated conductances and Na/K pump currents have been developed. This has revealed how Na/K pump currents contribute to rhythm and pattern generation in motor systems (Ellingson et al., 2021), intrinsic excitability in vertebrate neurons (Forrest et al., 2012), how pump currents mediate effects of neuromodulators (Forrest et al., 2012), and how pumps contribute to signal propagation through multi-compartment models (Scuri et al., 2007). Computational modeling in several neuronal cell types has also highlighted the importance of considering the dynamics of intracellular sodium concentration and its interactions with sodium-dependent pumps when examining neuronal dynamics, in particular on long time scales (Zylbertal et al., 2017a, Zylbertal et al., 2017b, Sharples et al., 2021, Kueh et al., 2016).

Multiple studies have anecdotally noted the presence of pump mediated effects on motor neuron excitability on multiple time scales, but exactly how specific pump parameters simultaneously shape intrinsic properties in motor neurons on both short (ms-s) and long (s-min) time scales remain relatively poorly understood. Paradoxically, Na/K pump activity is often described and modeled without the use of any explicit time constants. This raises the interesting question how long-lasting cellular events such as AHPs can be generated by instantaneous cellular processes. Overall, understanding how pump functional properties and the cellular ion concentration dynamics they contribute to shape excitability has relevance for human health, given that multiple human disorders are thought to arise from anomalous Na/K pump activity (reviewed in Holm and Lykke-Hartmann, 2016, and in Isaksen and Lykke-Hartmann, 2016).

Creating computational models of pump activity that can be deployed within the context of a genetically tractable model organism has value added, because the resulting models can then be coupled with genetic manipulations of cellular and circuit activity (Pulver and Griffith, 2010). Here we model Na/K pump activity in a Hodgkin-Huxley-type model based on a model of a motor neuron in *Drosophila* larvae, a system with an unsurpassed genetic toolkit. Our aim was to reveal conserved principles governing how modulation of specific Na/K pump functional properties shapes neuronal excitability on both short (ms-s) and longer (s-min) time scales. We deliberately chose to use a highly streamlined model with as few conductances as possible to maximize our ability to directly interpret effects of pump activity and generate a knowledgebase relevant across multiple species. We find that Na/K pumps have complex and often counterintuitive effects on intrinsic excitability, dynamics and cellular memory. Systematic exploration of the effects of modulating pump functional properties highlights the necessity of considering Na/K pump and Na reversal potential dynamics when interpreting the effects of experimental manipulations of cellular excitability and neural circuit function. This work provides insight and guidance for analogous work in vertebrate model organisms while also contributing to the groundwork for computational modelling in a genetically tractable invertebrate species.

## Methods and Materials

### Model neuron

We used a single compartment model neuron (Figure 1A) modified from (Gunay et al., 2015) for all simulations. In this model, membrane potential V is governed by:

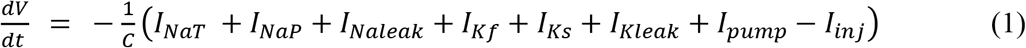

where C = 4.0 pF is the membrane capacitance, and the transient sodium current I_NaT_, the persistent sodium current I_NaP_, two potassium currents with fast (I_Kf_) and slow (I_Ks_) voltage-dependent gating dynamics, and sodium and potassium leak currents I_Naleak_ and I_Kleak_ are given by:

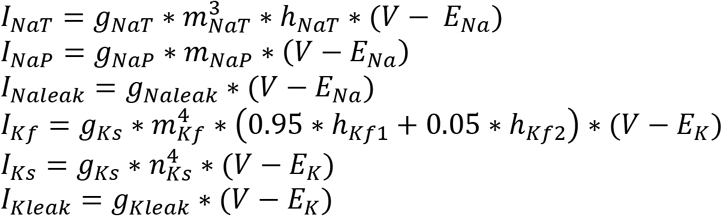

where g_NaT_ = 100.0 nS, g_NaP_ = 0.80 nS, g_Naleak_= 1.2 nS, g_Kf_ = 15.1 nS, g_Ks_ = 50.0 nS, g_Kleak_ = 3.75 nS are the maximal conductances for the respective currents. *E_Na_* is the sodium reversal potential (defined below), E_K_ = −80.0 mV is the potassium reversal potential, and the dynamic activation and inactivation variables are given by:

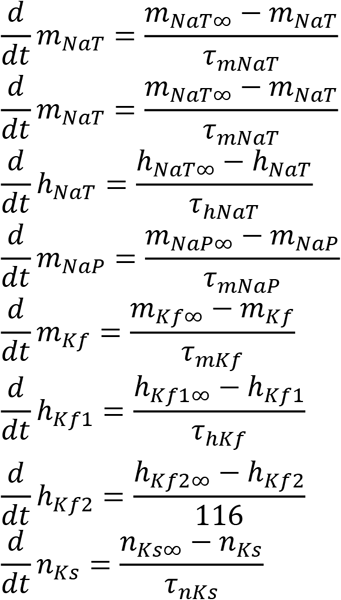

with voltage dependences and time constants

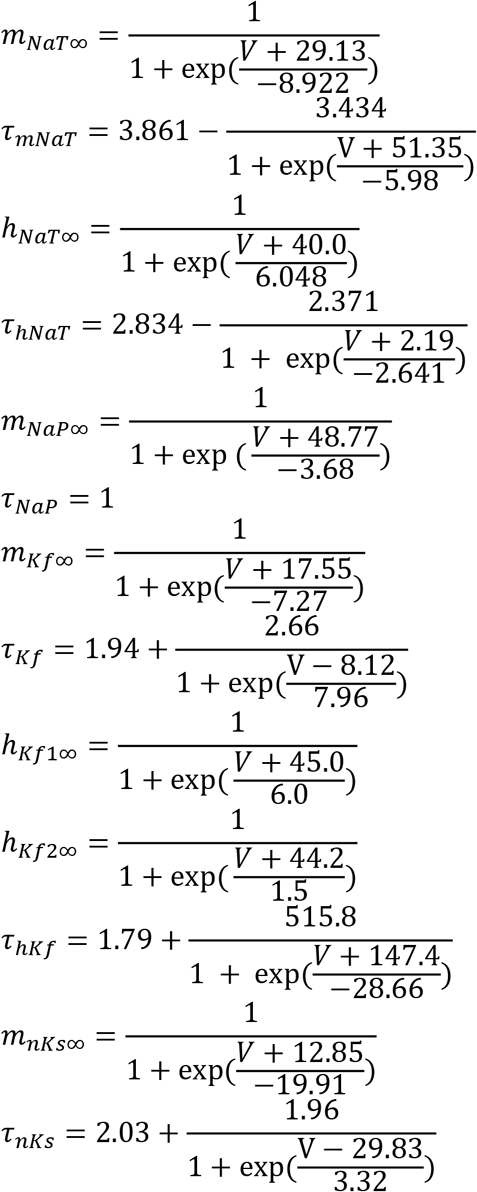

In the equations above, all times are in units of ms, all voltages are in units of mV.

**Figure 1.**
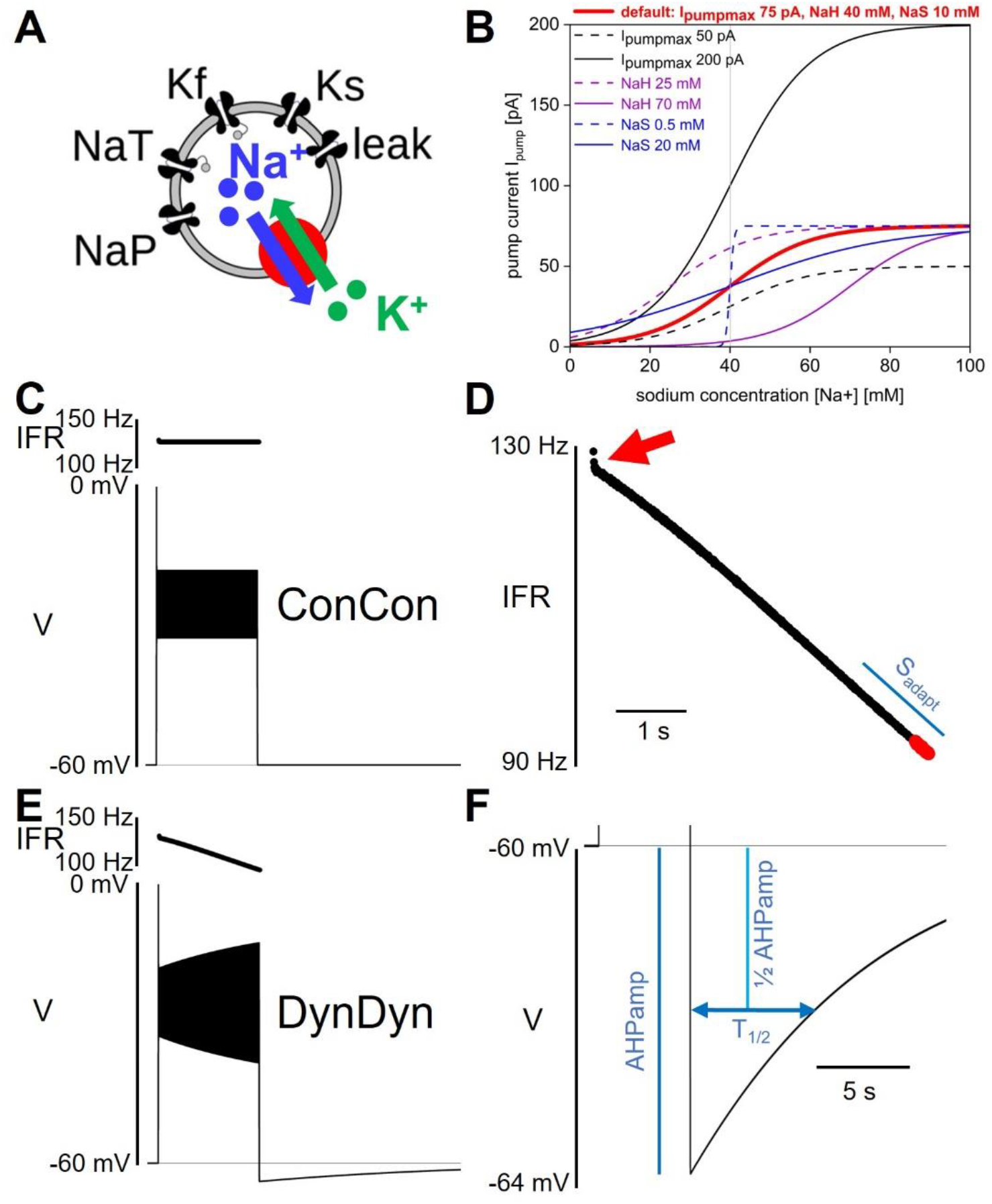
Model and pump properties, pump effects on spiking and after-hyperpolarization. (**A**) Schematic of model structure. (**B**) Dependence of Na/K pump current I_pump_ on intracellular sodium concentration [Na^+^] and pump parameters I_pumpmax_, NaH, and NaS. Red solid line shows pump activation curve with default pump parameters as listed at top, red. Other curves show pump activation curve for minimum (dashed) and maximum (solid) values of I_pumpmax_ (black), NaH (purple), and NaS (blue) explored in Figure 7. Grey vertical line indicates resting sodium concentration of model with default pump parameters. (**C)** Voltage response of model ConCon with constant [Na^+^] and E_Na_ to 5 s injection of 50 pA stimulus current. Insert above voltage trace shows instantaneous firing rate. **(D)** Expanded view of instantaneous firing rate IFR in DynDyn model, from E. Adaptation slope (s_adapt_) is indicated in blue and calculated based on trajectory of IFR in final periods of spiking (red dots, see Methods). Red arrow indicates initial rapid spike rate adaptation. (**E**) Same as C, but for model version DynDyn with dynamic [Na^+^] and E_Na_. Note constant IFR in ConCon, C, and adapting IFR in DynDyn, E. **(F)** Expanded view of DynDyn AHP, from E. Definitions of AHP amplitude (AHPamp) and AHP half duration (T_1/2_) are illustrated in blue.

The Na/K pump model is modified from (Kueh et al., 2016). The current I_pump_ is given by:

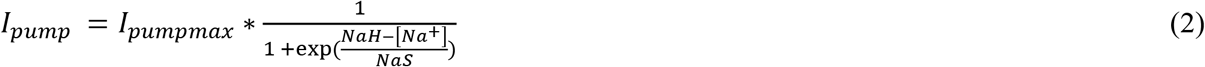

where I_pumpmax_ is the maximal pump current when the pump is fully activated, NaH is the sodium concentration at which the pump current is half-activated, and NaS is a factor that determines the steepness and range of the sodium concentration dependence of pump activation. Eq. 2 specifies a simple sigmoidal dependence of pump activation and pump current magnitude on intracellular sodium concentration as illustrated in Figure 1B (Kueh et al., 2016). Unless otherwise specified, the default values for the three pump parameters used here are ^I^pumpmax 75 pA, NaH = 40 mM, and NaS = 10 mM. Note that equation (2) does not contain an explicit time constant of pump activation, rather, pump current activation follows changes in [Na ^+^] instantaneously.

Together with the pump current I_pump_, the sodium currents I_NaT_, I_NaP_, and I_Naleak_ in this model alter the intracellular sodium concentration [Na^+^] via:

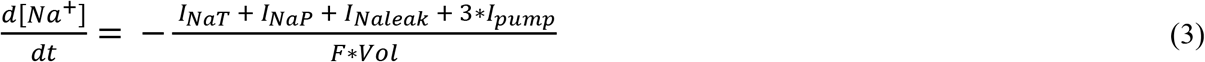

from its default value of [Na^+^] = 40.08 mM. Here, Vol = 549 μm^3^ is the volume of a shell under the neuronal membrane in which the sodium concentration is considered under the influence of sodium currents entering and exiting the cell, and the factor 3 multiplying the pump current accounts for the fact that each pump cycle extrudes three sodium ions and imports two potassium ions, corresponding to extrusion of three sodium ions for every net electrical charge change. Given the momentary intracellular sodium concentration [Na^+^] and a constant extracellular sodium concentration of 135 mM, the momentary sodium reversal potential E_Na_ driving Na^+^ fluxes through the membrane conductances is calculated according to the Nernst equation assuming a temperature of 25 °C.

To tease apart the roles of pump current, dynamic [Na^+^], and dynamic E_Na_ in neuronal excitability and memory, we introduced two switches in the model: a concentration switch that allows toggling the intracellular sodium concentration [Na^+^] between being constant at 40.08 mM versus being dynamic according to Eq. 3, and a reversal potential switch that allows toggling the sodium reversal potential E_Na_ between being constant at 31.2 mV (the Nernst potential corresponding to [Na^+^] = 40.08 mM) versus being dynamic according to the Nernst equation and the momentary value of [Na^+^]. These switches allow separating the direct effects of a time-varying intracellular sodium concentration from the indirect, electrical effects of the accompanying changes in sodium reversal potential E_Na_, which sets the driving force (V – E_Na_) for sodium flux through membrane channels. Where indicated in text and figures, model versions ConCon, DynCon, and DynDyn correspond to: version ConCon – [Na^+^] constant, E_Na_ constant; version DynCon – [Na^+^] dynamic, E_Na_ constant; and version DynDyn – [Na^+^] dynamic, E_Na_ dynamic.

### Measures of spiking activity and AHP features

We used several measures to quantify the spiking activity of model responses to step current injections of 5 s duration. Initial and final instantaneous firing rates IFR_ini_ and IFR_fin_ during the 5 s injection were calculated by taking the inverse of the first and last interspike intervals, respectively. Additionally, the rate of change in spike frequency was quantified as the slope s_adapt_ of the IFR profile at the end of the step current injection. Spike rate adaptation slope s_adapt_ was determined by taking the last 19 IFR values, separating them into two groups of 9, and averaging those two groups, while leaving off the very last IFR value (which in some simulations was invalid because the last spike was truncated by the step current injection ending mid-spike). The difference between the two IFR group averages, divided by the time difference between the middle spikes in each group of 9, is the adaptation slope s_adapt_ in units of Hz/s (see Figure 1D for illustration).

The after-hyperpolarization amplitude (AHPamp) was determined by taking the lowest voltage value after the end of the injection minus the pre-injection voltage. The after-hyperpolarization half duration (T_1/2_) was determined by finding the time after the end of the injection at which the AHP had decayed to 50% of the AHPamp value (see Figure 1F).

### Zap currents

The zap current injection was adopted from (Tohidi and Nadim, 2009). The zap current is given by:

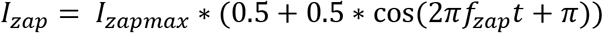

where I_zapmax_ is the maximum zap current amplitude (values used are specified in Figure 6), and f_zap_ is the exponentially increasing and decreasing zap frequency. For the acceleration portion of the zap, given f_zap_ is by:

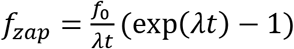

where f_0_ = 0.1 Hz is the minimum zap frequency, and t is time since the start of the acceleration portion of the zap injection. For the deceleration portion, the f_zap_ used is the mirror image (in time) of the acceleration f_zap_ function.

Lambda (λ) is the exponential rising factor given by:

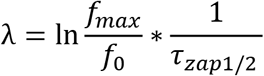

where f_max_ = 5.0 Hz is the maximum zap frequency, and *τ*_zap1/2_ = 20 s is half the total zap duration (acceleration and deceleration portions combined). The zap current generated by these equations is illustrated in Figure 6D.

### Model implementation and code availability

The model was implemented in Python (version 3.9). Differential equations were solved numerically using the fourth order Runge-Kutta algorithm ODEint available in the SciPy Python library. Voltage trace analysis such as spike detection, calculations of instantaneous firing rate, s_adapt_, measurement of AHPamp and T_1/2_, were implemented directly in the simulation code and executed at run-time. Simulation results were exported from the simulation code and imported into Origin 2022b (OriginLabs) for further analysis and plotting. All code is freely available on ModelDB at http://modeldb.yale.edu/267620.

## Results

### Na/K pump, dynamic sodium concentration and reversal potential added to basic model neuron

To examine the effects of Na/K pumps and intracellular sodium dynamics on the electrical activity and excitability of neurons, we incorporated a pump model and dynamic sodium concentration and reversal potential into a single-compartment, conductance-based neuron model. While the model neuron is loosely based on the electrical behavior of fly larval motoneurons and is a modified version of a previously published model, it does not include all the voltage-gated conductances known to be present in larval neurons (Lin et al., 2012, Gunay et al., 2015). Our intent was rather to study the effects of Na/K pumps and their parameters and dynamics using a basic pump model in a basic model neuron, so that we could assess the effects of modulating pump currents with as few confounds as possible and so that our findings may be applicable across a range of neuron types and neural systems that contain pumps. This approach complements studies on the other end of the scale of neuronal model complexity that examine the effects of pumps and sodium dynamics in multi-compartment models of specific neuron types with multiple membrane conductances and complex ion diffusion, buffering, and binding dynamics (Zylbertal et al., 2017a).

Briefly, our simplified model neuron contains two Na^+^ membrane currents, a transient sodium current I_NaT_ and a persistent sodium current I_Nap_, two K^+^ membrane currents with fast (I_Kf_) and slow (I_Ks_) voltage dependent gating dynamics, and a membrane leak current with a Na^+^ and a K^+^ component (Figure 1A). The equations governing the voltage-dependent dynamics of these currents and the model neuron’s membrane potential V are specified in the Methods section.

To this baseline model, we added a pump current I_pump_ whose activation depends on the intracellular sodium concentration [Na^+^] with the three parameters I_pumpmax_ (the maximal pump current when the pump is fully activated), NaH (the sodium concentration at which the pump current is half-activated), and NaS (a factor that determines the range and steepness of the sodium concentration dependence of pump activation) (Figure 1B). We further replaced the previously static intracellular sodium concentration [Na^+^] by a dynamic concentration variable influenced by sodium influx through the sodium membrane currents I_NaT_, I_NaP_, and I_Naleak_, and by sodium ion extrusion via the pump current. In contrast to more elaborate models of Na/K pumps and intracellular [Na^+^] dynamics (see for example, Zylbertal et al., 2017a), Eqs. (2) and (3), which describe pump activation and [Na^+^] dynamics, do not include any explicit activation time constants, buffering, or chemical reaction rate constants.

Addition of the pump current with default values of the three pump parameters of I_pumpmax_ = 75 pA, NaH = 40 mM, and NaR = 10 mM, combined with making the intracellular sodium concentration [Na^+^] and the sodium reversal potential E_Na_ dynamic, produced a silent (not spontaneously spiking or bursting) resting state of the model in which the pump is approximately half activated. This baseline activation can be thought of as the ‘housekeeping’ component of the pump’s activity, which balances the ongoing sodium influx in the resting state (primarily through the sodium component of the membrane leak conductance) to maintain a stable sodium concentration at rest. This particular resting state positions the [Na^+^]/E_Na_/pump dynamic system in the steepest part of the pump’s [Na^+^]-dependent activation curve with default pump parameters (bold red in Figure 1B), making it maximally sensitive to changes in [Na^+^] when the neuron is perturbed out of its resting state. However, our exploration of pump parameter space (see Figure 7, below) shows that our results are qualitatively similar and robust to pump parameter variations over wide ranges, and thus not idiosyncratic to this particular parameter combination and resting state.

Addition of the pump and of dynamic [Na^+^] and E_Na_ substantially altered the model’s spiking activity in response to step current injections (details below) and produced long-lasting (on the order of tens of seconds) afterhyperpolarizations (AHPs, Figure 1E). These AHPs show qualitative similarities to AHPs observed in larval *Drosophila* motor neurons and many other neuron types (Picton et al., 2017b).

### Effects of pump activity on spiking

We characterized how pump properties and dynamic [Na^+^] and E_Na_ affect spiking activity by applying a variety of stimulus types to the model neuron, the simplest (and most frequently used in electrophysiology experiments) being the injection of step currents. To quantify spiking activity during step current injections, we focused on three descriptive measures: the initial instantaneous firing rate IFR_ini_ at the beginning of step current injection, the final instantaneous firing rate IFR_fin_ at the end of a 5 s step current injection, and the slope of spike rate adaptation s_adapt_. How these measures are defined is described in the Methods section and in Figures 1D,F. In particular, we chose the slope s_adapt_ at the end of the instantaneous firing rate trajectory as a descriptor of the degree of firing rate adaptation because the instantaneous firing rate trajectories we observed in all model versions tended to have approximately linear shape over most of the 5 s current injection (see IFR profiles in Figures 1,2,3) that would be poorly characterized by other functions, such as an exponential fit.

**Figure 2.**
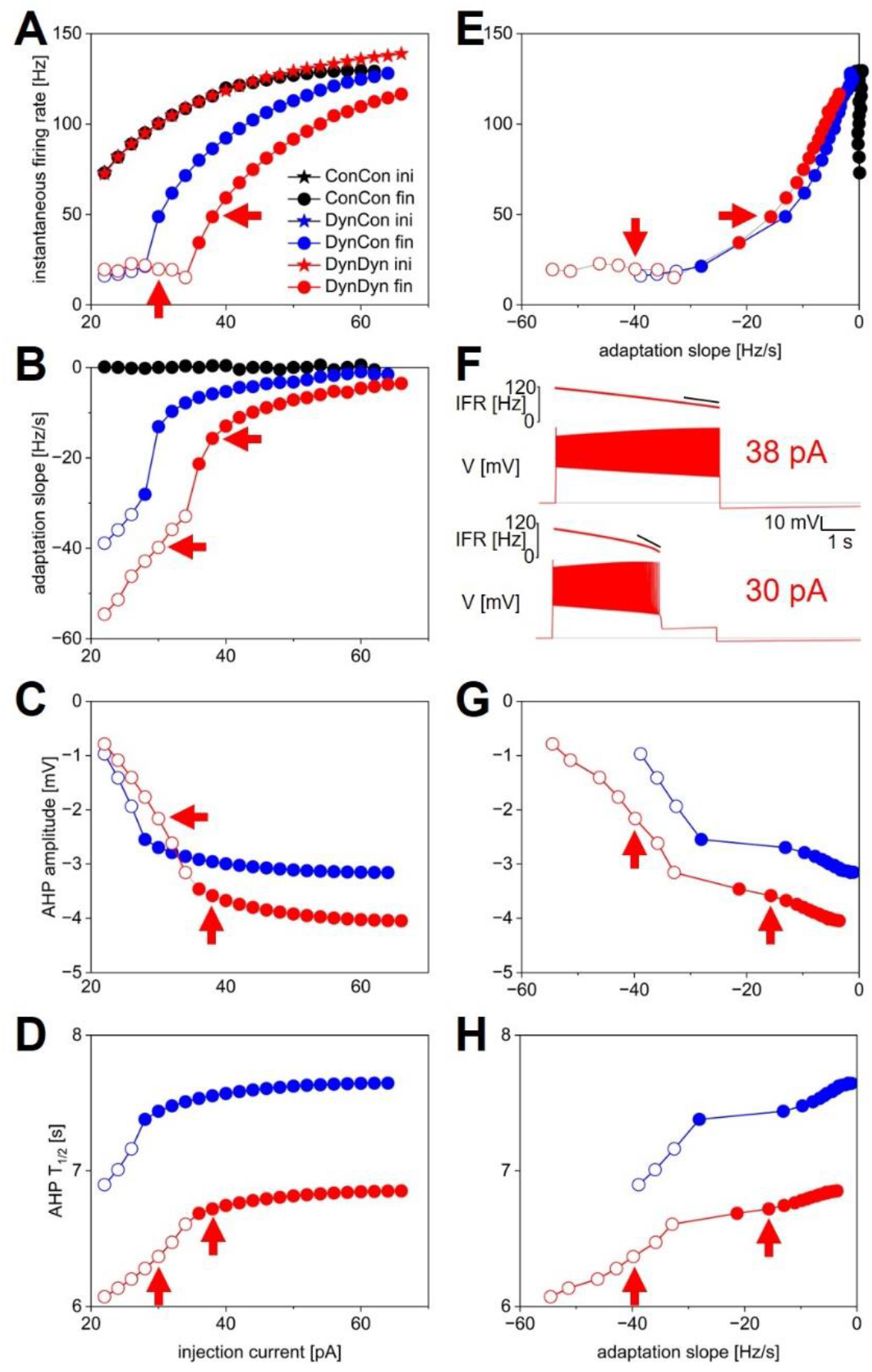
Measures of spiking activity and AHP dynamics in three model versions. **(A)** Initial (stars) and final (circles) instantaneous firing rate as a function of injection current in model versions with constant [Na^+^] and E_Na_ (ConCon, black), with dynamic [Na^+^] but constant E_Na_ (DynCon, blue), or with dynamic [Na^+^] and E_Na_ (DynDyn, red). Because initial IFR is almost identical in all three model versions, blue and black stars are occluded by red stars. Situations in which spiking stops before end of current injection in models DynCon and DynDyn are indicated by open circles. Red arrows indicate datapoints corresponding to traces in 2F. **(B)** Adaptation slope (sadapt) as a function of injection current. Color scheme same as in 2A, and throughout figure. **(C)** AHP amplitude as a function of injection current. **(D)** AHP half-duration T_1/2_ as a function of injection current. Model version ConCon omitted from 2C,D because of lack of AHP. **(E)** Same as 2A, but plotted against s_adapt_. **(F)** Example voltage traces showing full and incomplete spiking in DynDyn model for 38 pA (top) and 30 pA (bottom) injection currents, corresponding datapoints indicated by red arrows in all other panels. Black horizontal lines show pre-injection resting potential of −60mV. Instantaneous firing rate (IFR) is shown above voltage trace, with diagonal black lines indicating s_adapt_, which is −15.7 Hz/s for 38 pA injection (top) and −39.8 Hz/s for 30 pA injection (bottom). **(G)** Same as 2C, but plotted against s_adapt_. **(H)** Same as 2D, but plotted against s_adapt_.

**Figure 3.**
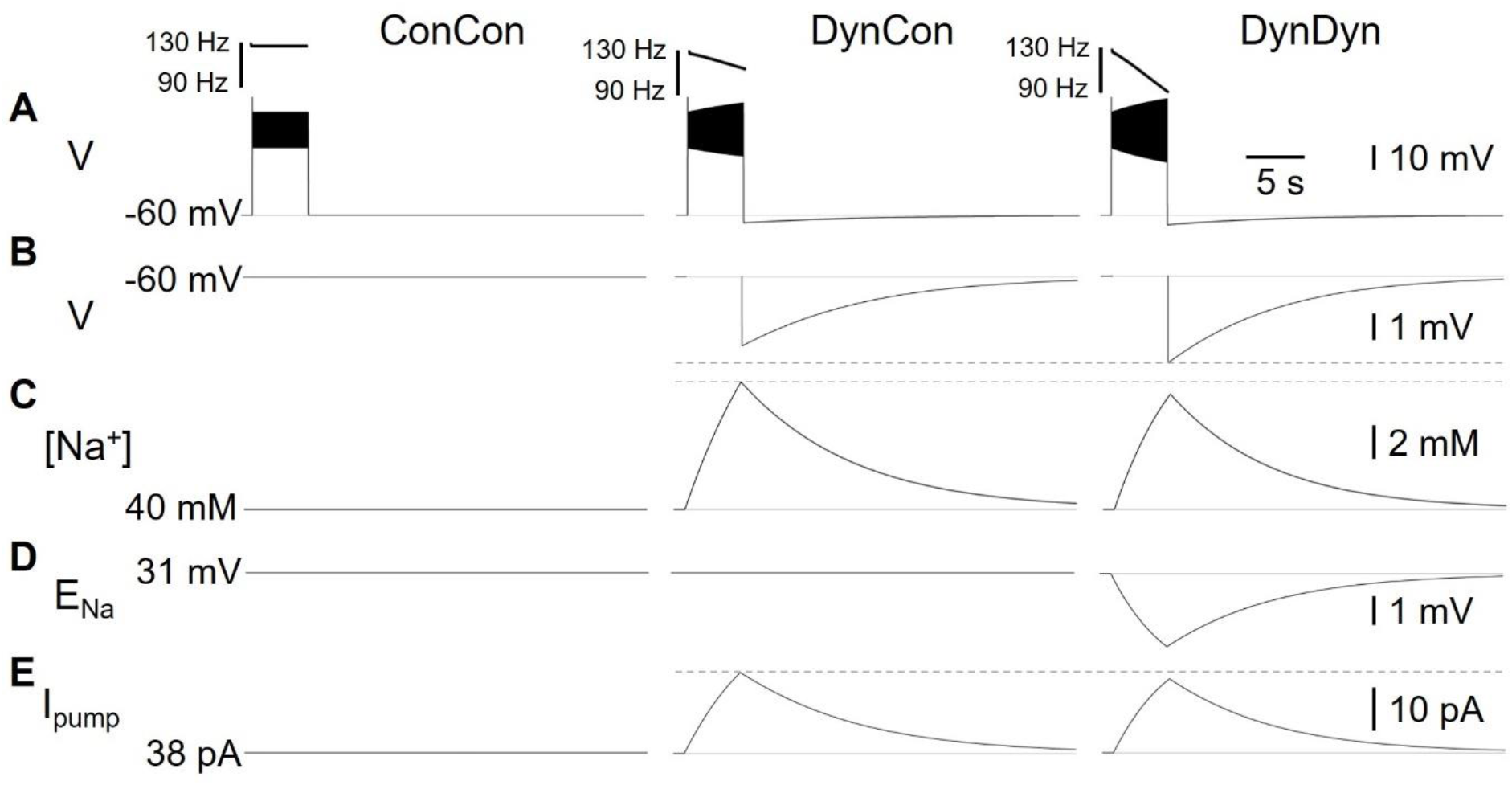
Effects of dynamic pump on spiking and AHP in model versions ConCon (left), DynCon (middle), and DynDyn (right). (**A**) Membrane potential V in response to 5 s, 50 pA current injection. Inserts above voltage traces show instantaneous firing rate IFR. (**B**) Membrane potential V at expanded vertical scale to show AHP. (**C**) Dynamics of intracellular sodium concentration [Na^+^] in the three model variants (**D**) Sodium reversal potential E_Na_. in all variants. (**E**) Pump current I_pump_ in variants. Dashed grey lines in B, C, E are inserted to facilitate comparison between DynCon and DynDyn.

Figure 2 quantifies these measures of spiking activity in response to step current injections of 5 s duration and of various amplitudes by comparing three model versions: version ConCon – which has constant values of [Na^+^] and E_Na_; version DynCon – in which [Na^+^] is dynamic, but E_Na_ is held constant; and version DynDyn – in which both [Na^+^] and E_Na_ are dynamic.

Note that because in living neurons, [Na^+^] and E_Na_ are directly linked through the Nernst equation, model version DynCon (in which [Na^+^] and E_Na_ are uncoupled) is an artificial construct, primarily intended to distinguish the effects of dynamic sodium concentration from the effects of dynamic changes in reversal potential (as we proceed to do in Figure 3) in a way that would not be possible in electrophysiology experiments, but is possible in a computational model. However, model version DynCon could also be interpreted as a simple, single-compartment proxy for a spatially extended neuron in which local changes in [Na^+^] that influence Na/K pumps in one location in the cell are spatially uncoupled from changes in E_Na_ that affect sodium influx through voltage-gated membrane channels in another location. Similarly, model version ConCon, in which neither [Na^+^] nor E_Na_ are allowed to vary, could be taken as a model of a patch-clamped neuron in which the intracellular ion concentrations are completely dominated by the large reservoir of patch pipette solution, and thus unable to vary. While spatial control of the intracellular milieu is rarely this complete in patch clamp experiments, considering this extreme case (implemented in ConCon) can be informative for the interpretation of experiments with partial control of the intracellular ion concentrations.

Note that comparison to a fourth model version in which the pump is entirely removed by setting I_pumpmax_ = 0 would not be physiologically meaningful in our simple model. In the absence of additional cellular mechanisms (Zylbertal et al., 2017a) that could help maintain sodium concentration, sodium reversal potential, and resting membrane potential, removing the pump entirely would lead to an unphysiological state in our simple model (see also Figure 7).

In all three model versions, ConCon, DynCon, and DynDyn, injection current amplitude was varied over a wide range, 0 pA to 100 pA, for Figure 2. For clarity, results from injection current levels that were either subthreshold (no spiking, at the low end of the injection current range) or produced depolarization block during part or all of the 5 s injection duration (at the high end of the injection current range) were omitted from Figure 2.

Initial firing rates at the beginning of step current injections of various amplitudes were virtually identical in model versions ConCon (black), DynCon (blue) and DynDyn (red), as Figure 2A indicates. This reflects that the resting values of [Na^+^] and E_Na_ in versions ConCon and DynCon were chosen to match the resting values that establish themselves in simulations of DynDyn after any numerical initialization artifact has decayed away. The three model versions thus are identical in their resting state – and therefore also in their firing rate immediately after the onset of current injection – and any differences only emerge after perturbation out of the resting state, in this case via step current injection.

In response to strong step current injection, spiking in all three model versions exhibits initial rapid spike rate adaptation over a few tens of ms (corresponding to a few spikes), as evident in the instantaneous firing rate profiles displayed in Figure 3A, and indicated in Figure 1D (red arrow). After this initial rapid adaptation, model version ConCon shows no further spike rate adaptation, whereas version DynCon shows intermediate and version DynDyn shows strong spike rate adaptation on the timescale of seconds (Figures 2A,B, inserts in 3A). Both versions with dynamic [Na^+^], DynCon and DynDyn, also exhibit a gradually increasing spike amplitude (in contrast to the constant spike amplitude in version ConCon).

While spiking in version DynDyn is continuous for the entire 5 s current injection when the injection current is above 35 pA, injection currents below 35 pA in version DynDyn produce spiking that ceases prematurely before the 5 s stimulus current ends (see Figure 2F, bottom for an example), because I_pump_ gradually increases during initial spiking activity, and eventually overwhelms the depolarizing injection current and inward membrane currents and suppresses further spiking despite ongoing stimulation. Similar premature termination of spiking also occurs in model DynCon, albeit at lower injection currents (below 28 pA, see open blue circles in Figure 2B). Notably, when spiking prematurely terminates in both DynCon and DynDyn, it does so once a similar final firing rate IFR_fin_ is reached, regardless of injection current – see open blue and red circles in Figure 2A, all are around 20 Hz. Interactions of pump currents with a dynamic intracellular sodium concentration and reversal potential thus appear to impose a limit on the maximal firing rate sustainable in a neuron, unless the injection current is so large that it dominates the membrane potential dynamics and forces the neuron to continue spiking despite high pump current activation. This effect of a dynamic pump/[Na^+^]/E_Na_ system could be interpreted as neuroprotective, in that it could potentially prevent overexcitation, exceedingly rapid spiking, and the accompanying risk of excitotoxity. Such a potentially protective effect is entirely absent in model version ConCon, where spiking at very high rates continues as long as a depolarizing injection current is provided (Figure 2A).

Figure 2 shows that addition of the Na/K pump with dynamic [Na^+^] and E_Na_ enriches neuronal spiking dynamics by adding slow spike rate adaptation and diversifying the nature of the spiking response to current steps. This addition transforms uniform spiking at a constant rate when the pump is merely a constant ‘housekeeper’ of sodium concentration, to a variable duration spiking response with spike rate adaptation when dynamic pump effects on sodium concentration and reversal potential are considered.

### Effects of pump on AHP features

Besides showing the effects of the pump and sodium concentration dynamics on spiking activity during step current stimulation, Figure 2 also illustrates dynamic pump effects on the AHP. We use two simple measures, the afterhyperpolarization amplitude (AHPamp) and half-duration (T_1/2_), as defined in the Methods section and illustrated in Figure 1F. While additional characterizations of the afterhyperpolarization, such as mathematical functions fitted to the AHP shape, could provide further information, AHPamp and T_1/2_ characterize basic features of AHP shape in condensed form.

Model version ConCon (black in Fig. 2) exhibits no AHP, whereas versions DynCon and DynDyn have long-lasting AHPs with membrane potential troughs dipping to −3.2 mV and −4.0 mV below resting membrane potential, respectively, for the larger depolarizing current injections (Figure 2C). These AHPs last for several tens of seconds, despite the absence of an explicit long time constant in the equations governing pump activation and sodium concentration dynamics (Eqs. 2 and 3 in Methods section). The AHP amplitude in model version DynDyn is larger than the AHP in version DynCon; this indicates that indirect electrical effects of pump-mediated sodium concentration changes via changes in sodium reversal potential E_Na_ can play an important role in how ion pumps shape cellular activity and memory.

The AHP in model DynDyn has a larger amplitude than that in DynCon (Figure 2C), but is also somewhat shorter, with half durations T_1/2_ in the range of 6 s to 7 s, compared to T_1/2_ values in the 7 s to 8 s range for version DynCon (Figure 2D). The AHP in DynDyn is therefore more ‘pointy’ (less shallow) than in DynCon.

### Relationship of AHP features to preceding spiking activity

The range of step current amplitudes explored in Figure 2 influences both features of the spiking response and features of the subsequent AHP. Previous experimental investigation of spiking, pump current, and AHP dynamics in fly larval motoneurons indicates that Na/K pump mediated AHPs can serve as spiking activity integrators and thus may constitute a form of cellular memory (Pulver and Griffith, 2010). To determine what kind and amount of information about preceding spiking might be contained in features of the AHP, we examined the relationships between spiking features IFRfin and s_adapt_, and AHP features AHPamp and T_1/2_ (Figures 2E,G,H). In the following description and analysis of this comparison, we focus on the range of injection current amplitudes for which spiking continues for the entire duration of the 5 s step current injection (filled circles in Figure 2E,G,H), to avoid the complexities inherent in lower level injection current simulations for which spiking terminates prematurely at different times for different levels of current injection (open circles in the same figures).

Interestingly, in both model versions DynCon (blue) and DynDyn (red), we found close to linear four-way relationships between IFR_fin_, s_adapt_, AHPamp, and T_1/2_, with more vigorous firing at stimulus offset and more pronounced spike rate adaptation corresponding to a deeper and longer-lasting AHP as the injection current was increased. In both models, features of the afterhyperpolarization therefore reflect information not only about the number of preceding spikes but also the dynamics of spiking. In contrast, the absence of an AHP in the ConCon model version means that just fractions of a second after vigorous spiking activity, any ‘memory’ of it in the form of persistent ion concentration changes and membrane hyperpolarization is lost if [Na^+^] and E_Na_ are not dynamic.

### Mechanisms underlying effects of pump activity

To investigate the electrical and Na^+^ concentration-related mechanisms producing and shaping the pump’s effects on spiking activity and longer time scale AHP features, we examined the time courses of multiple underlying dynamic variables in the model in addition to the membrane potential V, and for the same three model versions introduced in Figure 2. Figure 3 shows the dynamics of sodium concentration, reversal, and pump current in response to 5 s, 50 pA rectangular current pulses.

Intracellular [Na^+^] levels, while held constant in ConCon (Figure 3C, left), increase during spiking activity in DynCon (Figure 3C, middle) and DynDyn (Figure 3C, right) due to increased Na^+^ influx through the voltage-gated sodium channels underlying I_NaT_ and I_NaP_, which open during action potential firing. In both model versions, elevated [Na^+^] requires several tens of seconds to return back to baseline, far outlasting the spiking activity. The sodium concentration increase during stimulation is slightly smaller in DynDyn compared to DynCon. This is because the sodium driving force driving Na^+^ through the membrane conductances g_NaT_, g_NaP_, and g_Naleak_ is smaller in DynDyn, because in that model version, the sodium reversal potential E_Na_ is dynamic according to the Nernst equation, and is thus reduced during the [Na^+^] increase caused by spiking (Figure 3D).

Because the [Na^+^] increase is slightly smaller in DynDyn compared to DynCon, the pump is also slightly less activated, leading to a lower peak pump current at the end of spiking activity in DynDyn compared to DynCon (Figure 3E). Counter-intuitively, this smaller pump current nonetheless corresponds to a larger AHP amplitude in DynDyn compared to DynCon (−3.9 mV compared to −3.1 mV, Figure 3B). This is again explained by the reduced sodium driving force at the end of and immediately after current injection. The AHP is the result of a net hyperpolarizing current that in our simple neuron model arises from the balance of hyperpolarizing, outward currents I_Kf_, I_Ks_, I_Kleak_, and I_pump_, and depolarizing, inward currents I_NaT_, I_NaP_, and I_Naleak_, with the three latter smaller in DynDyn compared to DynCon because of the reduced value of E_Na_. The inward/outward current balance is therefore shifted more in favor of a net outward current in DynDyn compared to DynCon, which explains the deeper AHP.

Considering the inward/outward balance is also instructive when thinking about ion fluxes, not just electrical currents. The sodium membrane currents I_NaT_, I_NaP_, and I_Naleak_ provide avenues for sodium to flow into the cell and increase [Na^+^], which is counter-balanced by the pump reducing [Na^+^]. In our model, for default pump parameters as well as a wide range of other pump parameters, this balance is strongly in favor of sodium influx and [Na^+^] increase during spiking, when the voltage-gated channels underlying I_NaT_, I_NaP_ are open. In contrast, after spiking has ended and these channels close, the inward and outward sodium fluxes are nearly balanced, resulting in only a small net extrusion of sodium that takes tens of seconds to return the elevated intracellular sodium concentration to its baseline resting value (Figure 3C). This is the mechanism through which a [Na^+^]-dependent pump whose activation follows changes in [Na^+^] instantaneously, without an explicit time constant in Eq. (2), can nonetheless lead to a long-lasting memory trace in the form of elevated [Na^+^] and the accompanying AHP.

Although the differences in [Na^+^], E_Na_, AHP features, and Na/K pump dynamics between model versions DynCon and DynDyn appear modest in Figures 3B-E, the impact on spiking activity of holding the sodium reversal potential E_Na_ constant (model DynCon) as opposed to allowing it to vary with varying sodium concentration [Na^+^] (model DynDyn) is substantial. This is illustrated by the much stronger spike rate adaptation in DynDyn compared to DynCon (see steeper slope of IFR profile in Figure 3A, right compared to middle, and Figure 2A,B). It is therefore important to consider E_Na_ as a dynamic variable when modeling the role of Na/K pumps in neuronal activity, and when interpreting corresponding experimental data.

### Na/K pump effects on neuronal excitability and functional range: Test pulses

Figures 2 and 3 illustrate the effects of Na/K pumps and dynamic vs. constant [Na^+^] and E_Na_ on neuronal spiking activity and AHP features in response to step current injections. To investigate further how pumps affect neuronal excitability on multiple timescales, we simulated responses to several additional stimulation protocols designed to assess the extent to which the long-lasting dynamics of the [Na^+^]/E_Na_/I_pump_ system shape fast spiking, and vice versa.

In a first set of experiments, we assessed for how long pump mediated cellular memories persist. We delivered a short 200 ms, 22 pA test pulse to the model to probe its baseline excitability, then delivered a 50 pA, 5 s current stimulus to elicit spiking (Figure 4). Afterwards, at different time points during the course of the resulting AHP, we delivered the same short test pulse a second time in separate simulations that were executed independently, but that are overlaid in time in Figure 4B in different colors. For each test pulse individually, we examined the voltage response and measured the number of spikes fired during the 200 ms test pulse as a basic indicator of excitability (Figure 4D). Figure 4A shows traces from simulation runs in which the test pulse was delivered immediately after the AHP trough (1 s delay, red), immediately before and after the response transitioned from sub-threshold depolarization to spike firing (orange and green, 35 s and 36 s delays, respectively), and immediately before and after the response reached the same number of spikes (eight) as the pre-stimulus test pulse response (blue and black, 51 s and 52 s delays, with seven vs. eight spikes, respectively). Note that even when the number of spikes in response to the test current pulse has returned to the pre-stimulus number of eight, subtle differences remain between the voltage response at that time (52 s after the end of the main stimulus) and the pre-stimulus voltage response – compare for example the amplitudes of the last spikes in the two black traces in Figure 4A. It takes several more tens of seconds for the post-stimulus test pulse response to become indistinguishable from the pre-stimulus response. In addition, we confirmed that isolated test pulses given at the same extended time (e.g. 52 s) in the absence of the 5 s AHP generating stimulus elicited spiking responses that were identical to the initial response (data not shown). This indicates that pump activation and changes in [Na^+^] and E_Na_ during the spiking activity caused by the main 5 s stimulus affect cellular excitability even more than a minute after the end of the stimulation, at a time when the AHP (Figure 4B) and the accompanying change in intracellular sodium concentration (Figure 4C) appear at first glance to have returned to baseline, and in an electrophysiology experiment might be smaller than typical levels of recording noise.

**Figure 4.**
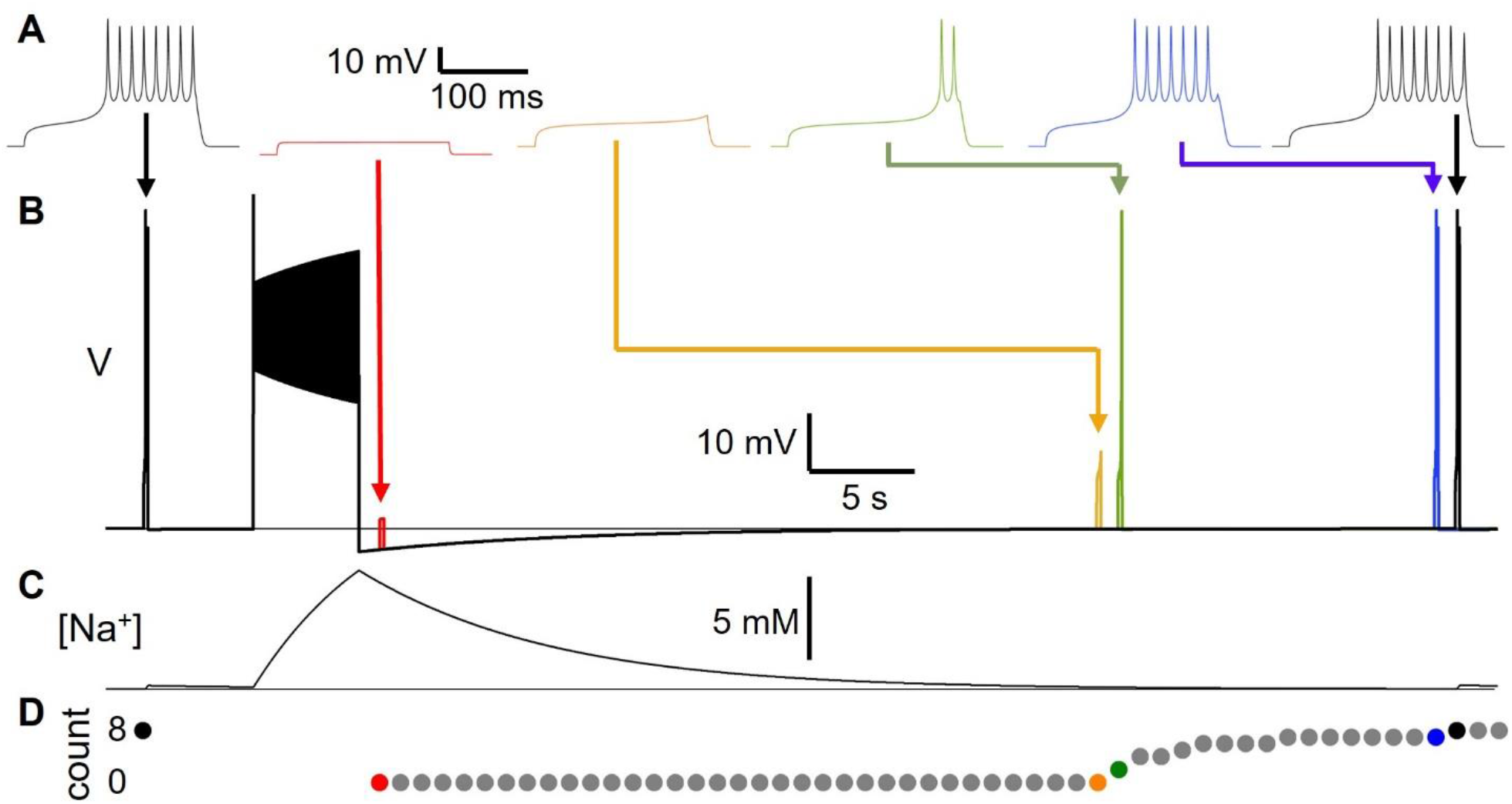
Long-lasting excitability changes following Na/K pump activation by spiking activity. (**A**) Voltage responses to 22 pA, 200 ms test pulses delivered to model version DynDyn – from left to right – before (black), and 1 s (red), 35 s (orange), 36 s (green), 51 s (blue) and 52 s (black) after a 50 pA, 5 s stimulus to generate spiking and a long lasting AHP. Arrows indicate timing of test pulses in B. (**B**) Superimposed simulations of individual test pulse responses and response to spike-inducing stimulus. Color code as in A. (**C**) Sodium concentration during simulation with test pulses delivered pre-stimulus and 52 s after stimulus, corresponding to black traces in A and B. (**D**) Number of spikes elicited by test pulses delivered before and once a second after main stimulus, in separate trials. Note that changes in excitability are present after AHP has finished and model has returned to original resting membrane potential.

### Na/K pump effects on neuronal excitability and functional range: Ramp currents

In a next set of experiments, we simulated ramps of injection current on different time scales, a stimulus protocol frequently used in electrophysiology to probe neuronal input-output relationships and dynamic properties. We simulated fast (2 s total duration) and slower (10 s total duration and 40 s total duration) bidirectional ramps of injection current from 0 pA to 70 pA, then back to 0 pA. The peak injected current of 70 pA was chosen as it is in the range of injection current for which model versions ConCon and DynDyn transition from spiking for the full duration of a 5 s step current injection, to depolarization block for part, or for all of the 5 s step current duration (Figure 2A).

Figure 5 shows the stimulus currents, voltage responses, and instantaneous firing rates as a function of momentary injection current for bidirectional ramps simulated in model versions ConCon (Figures 5A,D) and DynDyn (Figures 5B,E). The plots of instantaneous firing rate against momentary injection current in Figures 5D,E show clear firing hysteresis for slow and fast ramps in both model versions (i.e., with constant [Na^+^] and E_Na_ in ConCon, and dynamic [Na^+^] and E_Na_ in DynDyn). However, in ConCon spiking hysteresis only occurs in the form of spiking resuming at a lower injection current on the down ramp compared to the current at which spiking stopped on the up ramp, but not in terms of the instantaneous firing rate as a function of momentary injection current. In contrast, in the model version with dynamic [Na^+^] and E_Na_, IFR is consistently lower on the downward ramp compared to the upward ramp at the same momentary value of injection current, and this becomes more and more apparent with longer ramps. This is attributable to the activation of pump current as sodium accumulates intracellularly due to acceleration of spiking activity during the upward ramp; sodium levels then remain elevated and the pump current remains activated during the downward ramp, which sharply curtails spiking. Because the slower ramp stimuli allow more time for sodium to accumulate and the pump to be activated, the instantaneous firing rate at the same level of momentary injection current is consistently lower in the slowest ramp (blue in Figure 5E) than in the fastest ramp (red).

**Figure 5.**
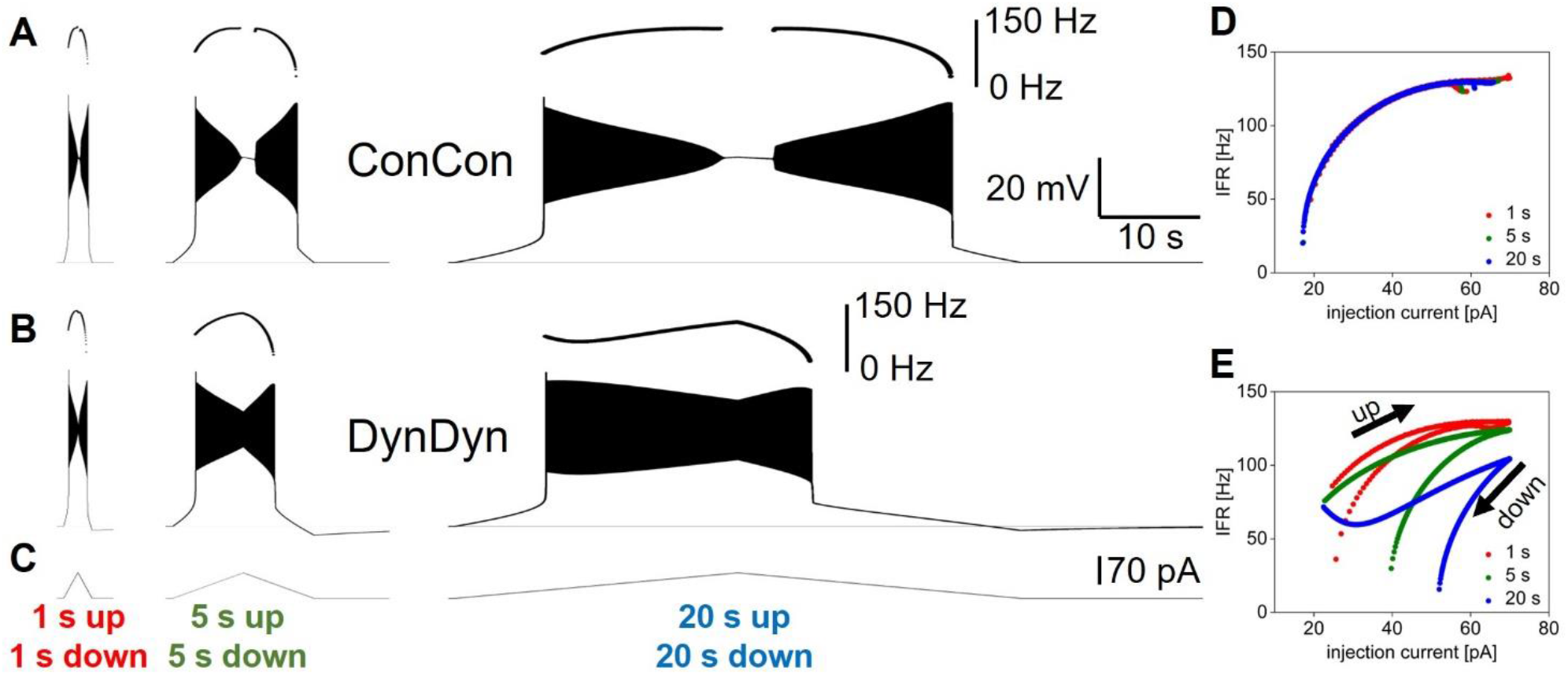
Influence of dynamic [Na^+^]/E_Na_/pump system on neuronal response to ramp currents. (**A**) Voltage response of model version ConCon to ramp currents with ramp up and ramp down durations of 1 s (left), 5 s (middle), and 20 s (right), and 70 pA peak amplitude. Inserts above voltage traces show instantaneous firing rate IFR. (**B**) Same for model version DynDyn. (**C**) Injected ramp currents. (**D**) Instantaneous firing rate from A (ConCon) as a function of momentary injection current for 1 s ramps (red), 5 s ramps (green), and 20 s ramps (blue). (**E**) Same for firing rates from B (DynDyn). Note similarities in response to fast ramps in A and B and increased divergence in IFR trajectory and hysteresis as duration of ramp increases.

**Figure 6.**
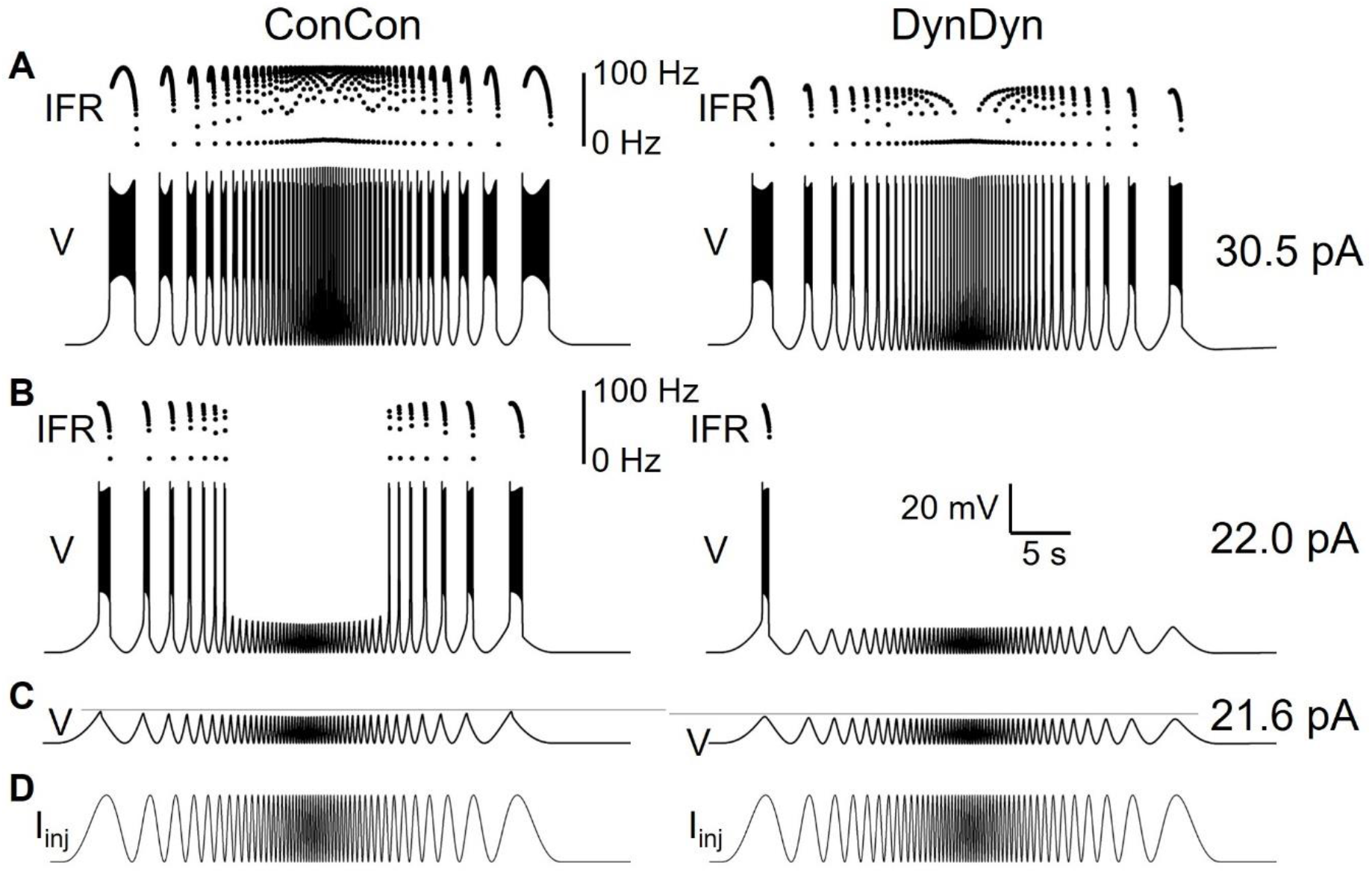
Dynamic [Na^+^]/E_Na_/I_pump_ system shapes neuronal response to oscillatory inputs of different frequencies. (**A**) Voltage response of model versions ConCon (left) and DynDyn (right) to a ‘zap’ injection current (described in Results section) of 30.5 pA amplitude and frequencies sweeping from 0.1 Hz to 5 Hz and back over a total duration of 40 s. Dots above voltage trace show instantaneous firing rate. (**B**) Same as A but for 22.0 pA zap current amplitude. (**C**) Voltage responses of ConCon (left) and DynDyn to 21.6 pA zap current. Horizontal grey lines included for ease of amplitude comparison, aligned to first voltage peak in each trace. Voltage and time scales in B (right) apply to A, B, and C. (**D**) Injected zap current time course showing increasing and decreasing stimulus frequency. Same time course was used in A, B, and C, but scaled vertically by current amplitudes given at right of A, B, and C, respectively.

**Figure 7.**
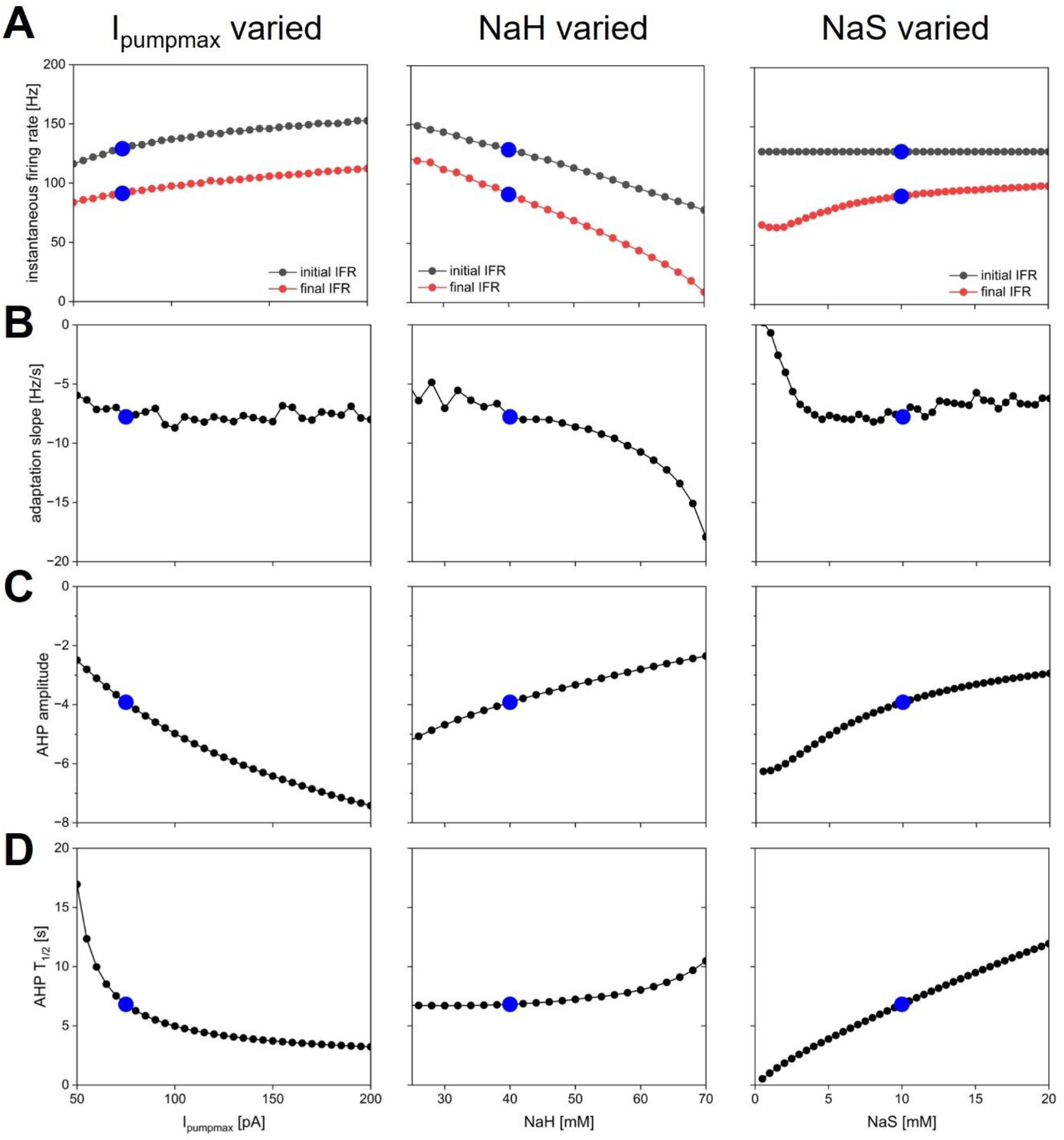
Dependence of spiking activity and AHP features on pump parameters. Each column shows variation of one pump parameter while keeping the other two at default value. Parameters varied were maximal pump current I_pumpmax_ (left column), sodium concentration NaH of pump half-activation (middle), and slope factor NaS of pump sodium-dependence (right). Blue dots show default value in each panel. (**A**) Initial (black) and final (red) instantaneous firing rate in response to a step current injection of 50 pA amplitude and 5 s duration (same stimulus as in Figure 3). (**B**) Spike rate adaptation slope s_adapt_ from same simulations. (**C**) AHP amplitude. (**D**) AHP half duration.

Notably, the presence of a dynamic [Na^+^]/E_Na_/pump system in the DynDyn model version reduced or prevented the depolarization block experienced by the ConCon version for injection currents near the peak of the bidirectional ramp, while also increasing the amount of current required to sustain spiking on the downward ramp, regardless of ramp speed (compare Figures 5A,D to B,E). Thus, sodium concentration and reversal potential dynamics altered the dynamic range of the model neuron, and ‘rescued’ it from depolarization block and spike amplitude attenuation, while making it less responsive to lower levels of injection current on the downward ramp.

Overall, Figure 5 shows that spiking dynamics in the DynDyn model are richer than in the ConCon model, with instantaneous firing rate reflecting not only the momentary level of injection current, but the previous history of injection current and spiking response experienced by the neuron. Notably, this richness is readily apparent with longer slower ramps, but less evident with shorter faster ramps. This finding could be taken into consideration when designing experimental stimulus protocols to probe slow dynamics in neuronal activity, in particular when the role of pumps is being investigated.

### Na/K pump effects on neuronal excitability and functional range: Zap currents

In a third set of experiments, we explored how pump currents and sodium and reversal dynamics shape responses to rhythmic current pulses of accelerating and decelerating frequency, termed ‘zap’ currents (Hutcheon and Yarom, 2000, Tseng and Nadim, 2010). This type of stimulus is highly relevant physiologically as larval motor neurons are typically recruited into rhythmic bursting through a range of frequencies (Pulver et al., 2015). We simulated sinusoidal, oscillating current injections that started with a 10 s cycle period (0.1 Hz frequency) and sped up to a cycle period of 200 ms (5 Hz frequency) over the course of 20 s with an exponential frequency profile (for details, see Methods section). Once cycle period reached 100 ms, the oscillations then returned to a 10 s cycle period over 20 s. Figure 6 shows the zap current (Figure 6D) and the corresponding voltage responses (Figures 6A,B,C) of model versions ConCon (left) and DynDyn (right).

A zap current of 30.5 pA amplitude evoked a supra-threshold voltage response in both model versions ConCon and DynDyn, with spikes occurring for every injected current peak, even up to the highest zap frequency, although at the fastest zap frequencies injected into DynDyn (middle of the voltage trace and IFR plot in Figure 6A, right) the model fired only a single spike per current peak, resulting in an instantaneous firing rate that tracked the zap frequency around its 5 Hz maximum in the middle of the zap injection. Consistent with the spike frequency-reducing effects of dynamic [Na^+^] and E_Na_ in Figures 2, 3, and 5, spike frequencies in response to zap current injection were also lower in DynDyn compared to ConCon throughout the zap simulation (see IFR plots in Figure 6A).

The overall activity profile of model version ConCon in both voltage and IFR is largely symmetric, showing very similar responses to the same zap frequencies in the two halves of the zap injection, whereas DynDyn responses to the same zap frequency in the deceleration half of the zap differed from the acceleration half – note for example the reduced burst duration in DynDyn in response to the last zap current peak compared to the first zap current peak. This indicates that with dynamic [Na^+^] and E_Na_ the neuron’s activity depends on its previous spiking history, whereas with constant [Na^+^] and E_Na_ it does not.

A zap current of 21.6 pA amplitude evoked a sub-threshold (non-spiking) voltage response in both model versions (Figure 6C), with relatively minor differences between the voltage responses. These differences primarily consisted of a slightly reduced voltage amplitude (first voltage peak of - 51.5 mV in DynDyn compared to −49.9 mV in ConCon, see horizontal grey lines in Figure 6C) and a very small (< 1 mV) AHP in DynDyn after the zap injection that is absent in ConCon.

Such subtle differences in sub-threshold voltage response would appear to have little effect on neuronal activity. However, the voltage responses to a zap current of 22.0 pA shown in Figure 6B reveal that even subthreshold voltage changes sculpted by a dynamic [Na^+^]/E_Na_/I_pump_ system can affect neuronal spiking response and endow a neuron with a long-lasting memory for not only supra-threshold, spiking activity, but also sub-threshold inputs and voltage fluctuations. This is evident in Figure 6B, right, where model version DynDyn responds to the first peak of the injected zap current with a brief burst of spikes, but fails to spike in response to the corresponding last current peak at the end of the zap injection. The subthreshold voltage fluctuations that occur between the first and last current peaks, by slightly activating the Na/K pump and changing [Na^+^] and E_Na_, have therefore left a memory trace that is reflected in DynDyn’s failure to spike in response to the last current peak.

Overall, our zap exploration of the effects of dynamic reversal and pump currents has revealed that pump currents shape how neurons respond to current injections on both fast and slow time scales.

### Systematic exploration of pump parameter space

Dynamic pump currents clearly generate a diversity of outputs in response to current stimuli. In Figures 1–6, we used a fixed set of pump parameters, I_pumpmax_ = 75 pA, NaH = 40 mM, NaR = 10 mM. How does modulating those parameters further shape spiking on short and long time scales in response to step current pulses? Our streamlined computational model enabled with relatively few cellular components allowed us to systematically vary all pump parameters and examine how each sculpts excitability. Figure 7 shows how key features of spiking and AHPs depend on I_pumpmax_, NaH and NaR. For each column in the figure, one of the parameters (left column – I_pumpmax_, middle – NaH, right – NaS) was varied over a physiologically meaningful range while the other two parameters were held at their default values. A step current of 50 pA amplitude and 5 s duration (same as in Figure 3) was simulated for each parameter combination, and spiking activity and AHP features were analyzed and plotted in Figure 7.

The ranges over which each parameter was varied were chosen as follows: I_pumpmax_ – lower bound set to 50 pA because I_pumpmax_ < 50 pA led to resting states with unphysiologically high [Na^+^] and low E_Na_; upper bound set to 200 pA because most effects of increasing I_pumpmax_ on spiking dynamics and AHP features appeared to saturate in that range. NaH – lower bound of 25 mM chosen because NaH < 25 mM led to depolarization block during part or all of the 5 s current injection; upper bound of NaH = 70 mM because NaH > 70 mM caused premature termination of spiking (as in Figure 2F, bottom) despite ongoing current injection. NaS – varied all the way down to 0.5 mM, which corresponds to an extremely steep pump current dependence on [Na^+^] (see dashed blue curve in Figure 1B), and up to 20 mM, where most effects of increasing NaS on spiking dynamics and AHP features appeared to saturate. The pump activation curves corresponding to the lower and upper bounds of these explored ranges are illustrated in Figure 1B in black for I_pumpmax_, purple for NaH, and blue for NaS.

Overall, the response of the model with dynamic [Na^+^] and E_Na_ (DynDyn) to step 5 s, 50 pA current injection did not qualitatively change over the parameter ranges covered in Figure 7, and consisted of spiking with spike rate adaptation for the duration of the stimulus injection, followed by a long-lasting (tens of seconds) AHP of several mV amplitude. Our model version DynDyn with our chosen default pump parameter set (indicated by blue dots in Figure 7) is therefore representative of a wide range of pump parameters.

The results in Figure 7 can be considered from two perspectives: 1) Which pump parameters primarily control a given feature of spiking or AHP? 2) Which spiking and AHP features does a given pump parameter control?

Figures 7A indicates that the instantaneous firing rate at the beginning and end of the 5 s current injection is influenced to a limited extent by the maximal pump current I_pumpmax_ and the slope factor NaS over most of their explored range, whereas the sodium concentration NaH of pump half-activation influences the firing rate more strongly, with higher values of NaH leading to lower spike rates. This seemingly counter-intuitive effect is explained by higher NaH leading to lower pump activation in the resting state, which results in higher resting [Na^+^] and correspondingly lower E_Na_, and an overall hyperpolarization of the model compared to the DynDyn model with default pump parameters. This more hyperpolarized state results in lower firing rates, all compared to default pump parameters.

The slope of spike rate adaptation s_adapt_ (Figure 7B) is also almost independent of all three pump parameters over a wide range, except for very large values of NaH (which produce steeply negative spike rate adaptation slopes), and very small values of NaS (which produce less spike rate adaptation). This latter effect occurs because the very steep dependence of pump activation on [Na^+^] for small values of NaS means that sodium influx due to a few spikes can rapidly activate the pump at the beginning of the 5 s injection, leaving little room for further pump activation and spike rate adaptation toward the end of the 5s injection, where s_adapt_ is measured.

The influence of all three pump parameters on AHP amplitude and half-duration T_1/2_ (Figure 7C,D) is monotonic and straightforward. AHPs are deeper (more negative values of AHPamp, Figure 7C) for larger I_pumpmax_ (because the pump is stronger), lower values of NaH (because the pump is activated at lower sodium concentrations), and lower values of NaS (because the pump activates over a narrower range of concentrations). In all three cases (three columns in Figure 7), larger AHP amplitudes (more negative values in Figure 7C) go along with shorter AHP half-durations (Figure 7D). This means that all three parameters can tune the AHP between a deep and pointy shape and a long and shallow shape, albeit the influence of NaH on T_1/2_ is relatively modest (Figure 7D, middle).

Taken together, our results illustrate that – at least in our simple neuron and pump model – pump parameters can vary widely while resulting in the same qualitative model behavior. Our results will therefore likely generalize to more complex pump and neuron models. Features of spiking and AHP, such as IFR, spike rate adaptation, and AHP amplitude, duration, and shape, are controlled by the interaction of all dynamic pump parameters, rather than one pump parameter controlling one activity or AHP feature independently. Our model therefore predicts that neuromodulation of pump currents or genetic alterations of pump proteins that affect the pump’s dependence on a dynamic intracellular sodium concentration can have complex and at times counter-intuitive effects that can be understood with the help of computational modeling.

### Modulation of pump properties shapes diversity of neuronal activity patterns

While manually sampling and exploring pump parameter space, we encountered a variety of pump parameter combinations that produced diverse activity patterns beyond our default model’s silence in the absence of current injection, and spiking response to 5 s current injection (Figures 2–4) or ramp or zap current injections (Figure 5). While a comprehensive exploration of the entire three-dimensional parameter space spanned by I_pumpmax_, NaH, and NaS is beyond the scope of this paper, Figure 8 illustrates examples of such different neuronal dynamics. Note that the examples in Figure 8 differ only in their three pump parameters, but have otherwise identical membrane conductance amplitudes and voltage-dependent dynamics, the same as our default DynDyn model.

**Figure 8.**
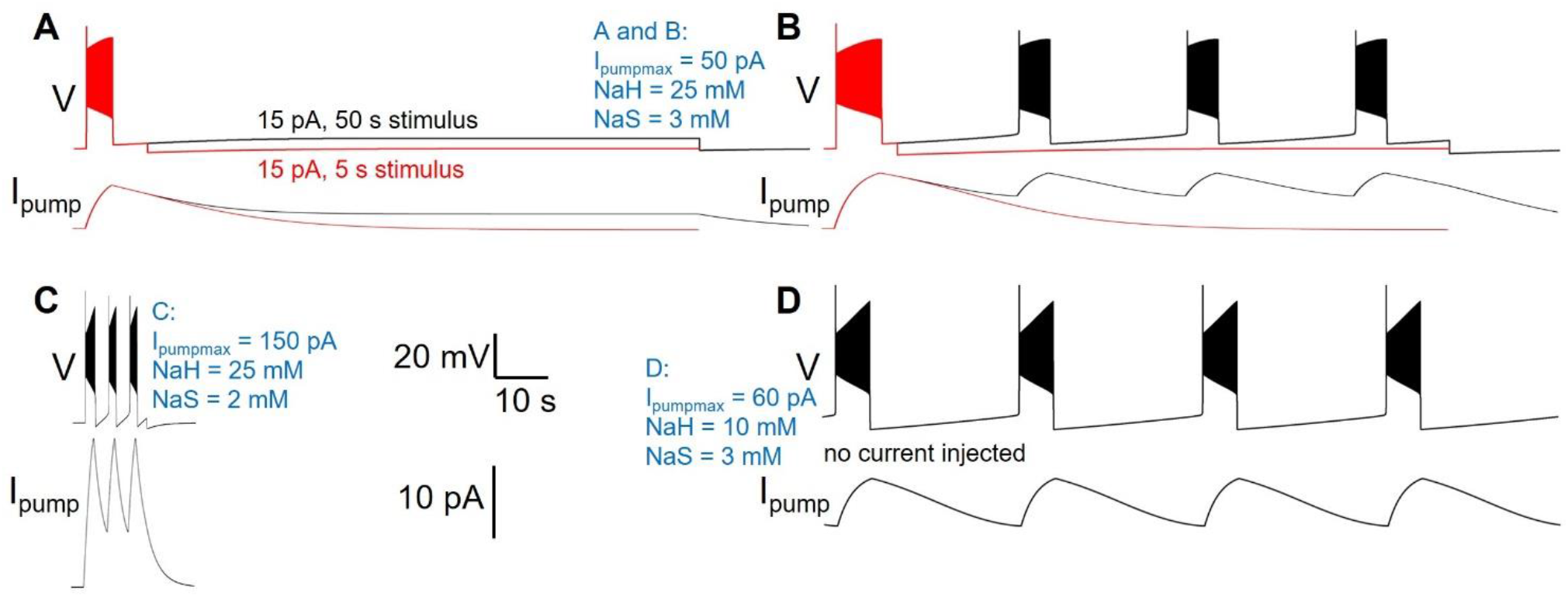
Different pump parameters produce a variety of activity types. Four examples of model version DynDyn with pump parameters varied (given in blue), but all other model parameters identical. (**A**) Pump parameters I_pumpmax_ = 50 pA, NaH = 25 mM, NaS = 3 mM produce premature termination of spiking in response to 15 pA, 5 s current injection (red). Spiking does not resume when 15 pA is continued for 50 s (black). Top: voltage; bottom: pump current. (**B**) In response to 20 pA, 5 s current injection, same pump parameters as in A also produce premature spike termination (red), but 50 s injection reveals repetitive bursting with 13.73 s burst period, 2.51 s burst duration, and 0.18 duty cycle (black). (**C**) Pump parameters I_pumpmax_ = 150 pA, NaH = 25 mM, NaS = 2 mM produce repetitive bursting with 1.69 s burst period, 0.57 s burst duration, and 0.33 duty cycle in response to 15 pA, 5 s current injection. (**D**) With pump parameters I_pumpmax_ = 60 pA, NaH = 10 mM, NaS = 3 mM, the model neuron becomes an endogenous burster in the absence of current injection, with 14.9 s burst period, 2.8 s burst duration, and 0.19 duty cycle. Same scale bars (bottom center of figure) apply to A – D.

Many pump parameter combinations produce continuous spiking in response to a step current injection of sufficient amplitude, as in Figures 2–4, but for a subset of pump parameter combinations we explored, spiking terminates before the end of 5 s stimuli of low current amplitude, as described above and shown in Figure 8A, where continuation of the stimulus current for 50 s reveals no further spiking, but a slightly depolarized silent state. In contrast, the same pump parameter combination (parameter values listed in Figure 8 caption, and in blue in the figure) enters a repetitive bursting regime when the injection current is slightly increased, from 15 pA to 20 pA (Figure 8B). We observed repetitive bursting in response to constant, small current injection for multiple pump parameter combinations, with burst periods, burst durations, and duty cycles (burst duration divided by burst period) covering wide ranges – Figure 8C shows an example of a pump parameter set that produces bursting on a much shorter time scale compared to that in Figure 8B. Finally, for some combinations of pump parameters, the DynDyn model neuron was transformed from a silent neuron in the absence of current injection, to an endogenously bursting neuron with oscillations of membrane potential and the [Na^+^]/E_Na_/I_pump_ system even without current injection (Figure 8D). A simple Na/K pump model in the context of a basic model neuron, as we explored here, can therefore endow the neuron with oscillatory properties on various time scales. This demonstrates that pump currents and dynamics of [Na^+^] and E_Na_ can substantially contribute to the richness of neuronal activity and response properties.

## Discussion

Here we explore how modulation of Na/K pump activity sculpts intrinsic properties in a conductance-based model of an invertebrate motor neuron. We find that long lasting AHPs following spike trains can be generated in a model neuron in which Na/K pump activity is modeled by a single type of Na/K pump that is dependent on a dynamically varying internal sodium concentration and a dynamically changing E_Na_. (model version DynDyn, Figure 3). AHPs can also be generated in models in which intracellular sodium is dynamic but E_Na_ is fixed (DynCon, Figure 3). We found that modelling E_Na_ as a dynamic variable can have profound effects on neuronal excitability. Strikingly, DynDyn models showed strong apparent spike frequency adaptation in response to current pulses. They also showed strong hysteresis in response to ramp up / ramp down stimuli, whereas ConCon models (with constant internal sodium and E_Na_) showed largely symmetric responses and very limited hysteresis (Figure 5). Similarly, DynDyn models showed asymmetric responses to oscillatory stimulation with oscillation frequencies accelerating and decelerating over a wide range (Figure 6). This suggests that dynamic E_Na_ could play a key supporting role in spike frequency adaptation and in encoding sodium pump mediated cellular memory in a variety of neuron types.

E_Na_ is widely considered to be a static parameter in neurons and is often modelled as such (Prinz et al., 2004, Gunay et al., 2015, Catterall et al., 2012). Traditional intracellular recording techniques by their nature tend to clamp or stabilize ionic concentrations inside and outside cells. In order to record from cells, it is necessary to expose them to relatively large volumes of physiological saline, and in some cases to largely dialyze out their internal solution, with patch pipette solutions. These recording conditions may impede a neuron’s activity to influence ionic concentrations sufficiently to affect E_Na_. In the animal however, where extracellular spaces are restricted by glial wrapping (Flanagan et al., 2018), cellular activity could conceivably shape ionic concentrations sufficiently to modulate E_Na_. Our modelling suggests that this could enrich cellular memory mechanisms. This raises the interesting (and unnerving) possibility that whole-cell recording techniques may be masking important and potentially highly conserved E_Na_ based intrinsic memory mechanisms in neurons. Hodgkin-Huxley type neuron models increasingly include simulations of intracellular and even extracellular ion concentrations, which enable researchers to explore how dynamically changing equilibrium potentials could shape excitability (Zylbertal et al., 2015). Non-invasive methods for optically measuring voltage could enable more detailed experiment study of processes that depend on dynamic E_Na_ and other equilibrium potentials (Miyazaki and Ross, 2015). However, voltage imaging methods may also increasingly generate results that conflict with or diverge from those obtained using traditional electrophysiological methods. Presence or absence of dynamic equilibrium potentials across preparations, and details of the recording technique used (sharp electrode vs whole cell vs perforated patch), could help explain these divergences.

Pump currents, together with the dynamics of intracellular sodium concentration and reversal potential, produce a long slow AHP that is evident in the membrane potential measured in electrophysiology experiments. Previous work has demonstrated how this long-lasting hyperpolarization allows cells to retain a memory of previous spiking through de-inactivation of voltage gated IA channels (Pulver and Griffith, 2010). But our results suggest that beyond what is typically considered the duration of the AHP – at a time after vigorous activity when the measured membrane potential has returned to within a fraction of one mV of the resting membrane potential – the consequence of a dynamic [Na^+^]/E_Na_/pump system can still silently shape neuronal excitability (Figure 4). These dynamics therefore create a window of ‘silent’ cellular memory for previous electrical activity that extends well beyond the experimentally obvious AHP. In our model, this memory mechanism arises solely from subtle lingering elevations of the sodium concentration that are only gradually returned to baseline because of the closely balanced interactions between inward sodium flux through membrane conductances, and outward sodium ion movement via the pump. This long-lasting biochemical integrator would be effectively invisible to noisy experimental voltage recording, but could shape responses to synaptic inputs in unpredictable ways. Optical imaging of sodium within neurons is a way to resolve whether these subtle events are present in living neurons, and explore how they interact with signaling molecules generated in neurons with more complex mixtures of ion channels and exchangers (Miyazaki and Ross, 2015, Meyer et al., 2022, Zylbertal et al., 2015).

The slow dynamics of [Na^+^] and E_Na_ influenced by Na/K pump action can furthermore shape the responses of a neuron to oscillatory inputs (Figure 6). Surprisingly, these dynamics can encode memories not just for prior spiking activity, but also for subthreshold membrane potential oscillations on a variety of time scales. Motor systems – particularly those of invertebrates – often operate on the time scale of seconds or tens of seconds and employ not only spike-mediated, but also graded synaptic transmission, in which transmitter release is dependent on subthreshold membrane potential fluctuations (Prinz et al., 2003, Ivanov and Calabrese, 2006a, Ivanov and Calabrese, 2006b). Previous work has characterized how pump currents can mediate memory of spiking and how neuromodulators sculpt those memories (Hachoumi et al., 2022). Our results suggest that pump-based cellular memory could have further profound effects on motor systems because it can shape motor neuron recruitment patterns in a manner that is dependent on both prior spiking *and subthreshold* activity of the motor system, as well as its neuromodulatory state.

Pump properties can also generate what appears at first to be a relatively simple spike frequency adaptation; but upon further investigation it is actually a manifestation of a slow bursting mechanism (Figure 8). Whether or not this bursting mechanism is apparent on any given time scale will be determined in part by the combination of sodium pump parameters in a given cell and the experimental protocols used for measuring excitability. Previous work has demonstrated how Na/K pumps can interact with hyperpolarization-activated currents Ih to generate episodic bursting in vertebrate spinal cord (Sharples et al., 2021) and leech heartbeat generator (Kueh et al., 2016). Extensive work in cardiac physiology has also demonstrated a key role for Na/K pumps in myogenic rhythms (reviewed in McDonough et al., 2002). Our results demonstrate that a simpler bursting mechanism is also possible in which depolarizations lead to bursts of spikes that trigger sodium influx. This then shifts E_Na_ and activates an outward current which hyperpolarizes neurons, and restarts a cycle. Similar bursting and oscillations with involvement of the Na/K pump current have previously been described in another simple Hodgkin-Huxley-type model and analyzed using a dynamical systems approach to separate the fast (spiking) time scale from a slow time scale arising from pump contributions to ion concentration changes (Barreto and Cressman, 2011). Bursting with pump involvement has further been noted in more complex individual neurons and in small circuits of more complex neurons with pumps and additional synaptic dynamics (Ellingson et al., 2021), or as arising from the interaction of pump-supported intrinsic dynamics and network connectivity (Zylbertal et al., 2017b). In our model, bursting occurs in the absence of membrane currents that typically support bursting dynamics, such as the hyperpolarization-activated current Ih, or oscillations on the basis of calcium currents and the calcium-dependent potassium current I_KCa_. Pump-supported oscillation cycles can be remarkably long (i.e. tens of seconds, Figure 8) and therefore could be playing important evolutionarily conserved roles in generating oscillations underlying longer time scale rhythmic activities like sleep (Tabuchi et al., 2018) and mating (Wagenaar et al., 2010). Indeed sodium dependent mechanisms are thought to contribute to ‘infra-slow’ oscillations in vertebrate olfactory neurons (Zylbertal et al., 2017b) (Wagenaar et al., 2010).

Our exploration of pump parameter space suggests that in addition to affecting the responsiveness of a neuron to inputs, pump properties can also qualitatively change the spontaneous activity produced by a neuron. This includes the transformation of silent neurons (that spike or burst only in response to inputs), into endogenously spiking or bursting neurons via moderate changes to the dependence of pump activation on the intracellular sodium concentration (Figure 8). Many invertebrate motor circuits are composed of both types of neurons, silent ‘follower’ neurons as well as endogenously oscillating ‘pacemaker’ neurons (reviewed in Marder and Calabrese, 1996, Marder et al., 2022). Furthermore, the same neurons that act as motor neurons and control muscles can also act as interneurons and participate in pattern generation based on their oscillatory and response properties. Modifying pump properties that affect these neurons’ excitability and responsiveness, for example through neuromodulation, could therefore provide mechanisms to sculpt rhythm generation with pumps playing roles that go beyond simple cellular housekeeping.

Studies of invertebrate neural circuits have consistently demonstrated a core principle: To understand how a circuit generates outputs, it is critical to measure the dynamics of synaptic transmission and the dynamic intrinsic properties of individual neurons within a circuit. The dynamics of synaptic transmission and voltage-gated ionic conductances in identified neurons within circuits have been well explored in multiple systems (reviewed in Marder and Calabrese, 1996, Marder et al., 2022). However, the dynamics of Na/K pump currents are less well understood. Given that Na/K pumps are highly conserved across all animal phyla, and play fundamental roles in shaping excitability, it makes sense to establish knowledgebases for understanding how dynamics of pump activity influence intrinsic properties of neurons, which in turn, shape neural circuit activity. Genetically tractable invertebrate nervous systems such as the *Drosophila* larval locomotor system, in combination with computational modelling, provide attractive vehicles for crawling into this space.

## Acknowledgements

We thank Ben Orli-Nathanson, Julius Jonaitis, and Lindsay Hexter for their valuable contributions during the initial phases of this project. We would like to thank Eve Marder, Leslie Griffith, Don Katz and the Brandeis Neuroscience Program for supporting inspirational work on invertebrate and vertebrate neural circuits, and for the Brandeis community’s steadfast commitment to training and mentoring young scientists over many decades.

## References

Barreto, E. & Cressman, J. R. 2011. Ion concentration dynamics as a mechanism for neuronal bursting. J Biol Phys, 37, 361–73.

Catterall, W. A., Raman, I. M., Robinson, H. P., Sejnowski, T. J. & Paulsen, O. 2012. The Hodgkin-Huxley heritage: from channels to circuits. J Neurosci, 32, 14064–73.

Ellingson, P. J., Barnett, W. H., Kueh, D., Vargas, A., Calabrese, R. L. & Cymbalyuk, G. S. 2021. Comodulation of h-and Na(+)/K(+) Pump Currents Expands the Range of Functional Bursting in a Central Pattern Generator by Navigating between Dysfunctional Regimes. J Neurosci, 41, 6468–6483.

Flanagan, B., Mcdaid, L., Wade, J., Wong-Lin, K. & Harkin, J. 2018. A computational study of astrocytic glutamate influence on post-synaptic neuronal excitability. PLoS Comput Biol, 14, e1006040.

Forrest, M. D., Wall, M. J., Press, D. A. & Feng, J. 2012. The sodium-potassium pump controls the intrinsic firing of the cerebellar Purkinje neuron. PLoS One, 7, e51169.

Gunay, C., Sieling, F. H., Dharmar, L., Lin, W. H., Wolfram, V., Marley, R., Baines, R. A. & Prinz, A. A. 2015. Distal spike initiation zone location estimation by morphological simulation of ionic current filtering demonstrated in a novel model of an identified Drosophila motoneuron. PLoS Comput Biol, 11, e1004189.

Hachoumi, L., Rensner, R., Richmond, C., Picton, L., Zhang, H. & Sillar, K. T. 2022. Bimodal modulation of short-term motor memory via dynamic sodium pumps in a vertebrate spinal cord. Curr Biol, 32, 1038–1048 e2.

Holm, T. H. & Lykke-Hartmann, K. 2016. Insights into the Pathology of the α3 Na+/K+-ATPase Ion Pump in Neurological Disorders; Lessons from Animal Models. Frontiers in Physiology, 7.

Hutcheon, B. & Yarom, Y. 2000. Resonance, oscillation and the intrinsic frequency preferences of neurons. Trends Neurosci, 23, 216–22.

Isaksen, T. J. & Lykke-Hartmann, K. 2016. Insights into the Pathology of the α2-Na+/K+-ATPase in Neurological Disorders; Lessons from Animal Models. Frontiers in Physiology, 7.

Ivanov, A. I. & Calabrese, R. L. 2006a. Graded inhibitory synaptic transmission between leech interneurons: assessing the roles of two kinetically distinct low-threshold Ca currents. J Neurophysiol, 96, 218–34.

Ivanov, A. I. & Calabrese, R. L. 2006b. Spike-mediated and graded inhibitory synaptic transmission between leech interneurons: evidence for shared release sites. J Neurophysiol, 96, 235–51.

Kaplan, J. H. 2002. Biochemistry of Na,K-ATPase. Annu Rev Biochem, 71, 511–35.

Kueh, D., Barnett, W. H., Cymbalyuk, G. S. & Calabrese, R. L. 2016. Na(+)/K(+) pump interacts with the h-current to control bursting activity in central pattern generator neurons of leeches. Elife, 5.

Lin, W. H., Gunay, C., Marley, R., Prinz, A. A. & Baines, R. A. 2012. Activity-Dependent Alternative Splicing Increases Persistent Sodium Current and Promotes Seizure. Journal of Neuroscience, 32, 7267–7277.

Marder, E. & Calabrese, R. L. 1996. Principles of rhythmic motor pattern generation. Physiol Rev, 76, 687–717.

Marder, E., Kedia, S. & Morozova, E. O. 2022. New insights from small rhythmic circuits. Curr Opin Neurobiol, 76, 102610.

Mcdonough, A. A., Velotta, J. B., Schwinger, R. H., Philipson, K. D. & Farley, R. A. 2002. The cardiac sodium pump: structure and function. Basic Res Cardiol, 97 Suppl 1, I19–24.

Meyer, D. J., Diaz-Garcia, C. M., Nathwani, N., Rahman, M. & Yellen, G. 2022. The Na(+)/K(+) pump dominates control of glycolysis in hippocampal dentate granule cells. Elife, 11.

Miyazaki, K. & Ross, W. N. 2015. Simultaneous Sodium and Calcium Imaging from Dendrites and Axons. eNeuro, 2.

Picton, L. D., Nascimento, F., Broadhead, M. J., Sillar, K. T. & Miles, G. B. 2017a. Sodium Pumps Mediate Activity-Dependent Changes in Mammalian Motor Networks. J Neurosci, 37, 906–921.

Picton, L. D., Sillar, K. T. & Zhang, H. Y. 2018. Control of Xenopus Tadpole Locomotion via Selective Expression of Ih in Excitatory Interneurons. Curr Biol, 28, 3911–3923 e2.

Picton, L. D., Zhang, H. & Sillar, K. T. 2017b. Sodium pump regulation of locomotor control circuits. J Neurophysiol, 118, 1070–1081.

Prinz, A. A., Bucher, D. & Marder, E. 2004. Similar network activity from disparate circuit parameters. Nat Neurosci, 7, 1345–52.

Prinz, A. A., Thirumalai, V. & Marder, E. 2003. The functional consequences of changes in the strength and duration of synaptic inputs to oscillatory neurons. J Neurosci, 23, 943–54.

Pulver, S. R., Bayley, T. G., Taylor, A. L., Berni, J., Bate, M. & Hedwig, B. 2015. Imaging fictive locomotor patterns in larval Drosophila. J Neurophysiol, 114, 2564–77.

Pulver, S. R. & Griffith, L. C. 2010. Spike integration and cellular memory in a rhythmic network from Na+/K+ pump current dynamics. Nat Neurosci, 13, 53–9.

Scuri, R., Lombardo, P., Cataldo, E., Ristori, C. & Brunelli, M. 2007. Inhibition of Na+/K+ ATPase potentiates synaptic transmission in tactile sensory neurons of the leech. Eur J Neurosci, 25, 159–67.

Sharples, S. A., Parker, J., Vargas, A., Milla-Cruz, J. J., Lognon, A. P., Cheng, N., Young, L., Shonak, A., Cymbalyuk, G. S. & Whelan, P. J. 2021. Contributions of h-and Na(+)/K(+) Pump Currents to the Generation of Episodic and Continuous Rhythmic Activities. Front Cell Neurosci, 15, 715427.

Tabuchi, M., Monaco, J. D., Duan, G., Bell, B., Liu, S., Liu, Q., Zhang, K. & Wu, M. N. 2018. Clock-Generated Temporal Codes Determine Synaptic Plasticity to Control Sleep. Cell, 175, 1213–1227 e18.

Tohidi, V. & Nadim, F. 2009. Membrane resonance in bursting pacemaker neurons of an oscillatory network is correlated with network frequency. J Neurosci, 29, 6427–35.

Tseng, H. A. & Nadim, F. 2010. The membrane potential waveform of bursting pacemaker neurons is a predictor of their preferred frequency and the network cycle frequency. J Neurosci, 30, 10809–19.

Wagenaar, D. A., Hamilton, M. S., Huang, T., Kristan, W. B. & French, K. A. 2010. A hormone-activated central pattern generator for courtship. Curr Biol, 20, 487–95.

Zhang, H. Y. & Sillar, K. T. 2012. Short-term memory of motor network performance via activity-dependent potentiation of Na+/K+ pump function. Curr Biol, 22, 526–31.

Zylbertal, A., Kahan, A., Ben-Shaul, Y., Yarom, Y. & Wagner, S. 2015. Prolonged Intracellular Na+ Dynamics Govern Electrical Activity in Accessory Olfactory Bulb Mitral Cells. PLoS Biol, 13, e1002319.

Zylbertal, A., Yarom, Y. & Wagner, S. 2017a. The Slow Dynamics of Intracellular Sodium Concentration Increase the Time Window of Neuronal Integration: A Simulation Study. Front Comput Neurosci, 11, 85.

Zylbertal, A., Yarom, Y. & Wagner, S. 2017b. Synchronous Infra-Slow Bursting in the Mouse Accessory Olfactory Bulb Emerge from Interplay between Intrinsic Neuronal Dynamics and Network Connectivity. J Neurosci, 37, 2656–2672.

